# Neurophysiological correlates of residual inhibition in tinnitus: Hints for trait-like EEG power spectra

**DOI:** 10.1101/2020.12.15.422892

**Authors:** S. Schoisswohl, M. Schecklmann, B. Langguth, W. Schlee, P. Neff

## Abstract

Magneto- and electroencephalography (M/EEG) investigations in tinnitus patients demonstrated anomalous oscillatory brain activity patterns compared to healthy controls. A well-established phenomenon in tinnitus is the possibility to temporary suppress tinnitus following acoustic stimulation, which is termed residual inhibition (RI). The few former neurophysiological investigations of RI reported partly conflicting results hampering consensus on tinnitus-specific brain activity and basic neural models.

Hence, our objective was to investigate RI-specific oscillatory brain activity changes and whether these changes can be associated with behavioral measures of tinnitus loudness. Further, contrasts between acoustic stimulation responders and non-responders provide further insights in RI-related spontaneous brain activity.

Three different types of noise stimuli were administered for acoustic stimulation in 45 tinnitus patients. Subjects resting state brain activity was recorded before and during RI via EEG alongside with subjective measurements of tinnitus loudness.

On the whole-group level, tinnitus-unspecific changes were observed which fit established knowledge about basic neural responses after acoustic stimulation. Responder non-responder contrasts revealed differences in alpha and gamma band activity in line with the proposed neural models for oscillatory brain activity in tinnitus. Further analysis of sample characteristics demonstrated divergences between responders and nonresponders notably for tinnitus duration. During RI, distinct differences between responders and non-responders were exclusively observed for alpha band activity in auditory cortical areas. Neither correlations of behavioral tinnitus measures nor differences between stimulus-induced changes in ongoing brain activity could be detected.

Taken together, our observations might be indicative of trait-specific forms of oscillatory signatures in different subsets and chronification grades of the tinnitus population possibly related to acoustic tinnitus suppression. Results and insights are not only useful to understand basic neural mechanisms behind RI but are also valuable for general neural models of tinnitus.

**Highlights:**

- Residual inhibition provides a key method to study the basic mechanisms of tinnitus.
- We compared residual inhibition EEG activity between responders and non-responders.
- In responders, the alpha activity in auditory areas was increased during tinnitus suppression.
- Results and insights are valuable for understanding the neural mechanisms behind acoustic tinnitus suppression.

## 1 Introduction

Subjective tinnitus is defined as the perception of a ringing or hissing without the presence of a corresponding internal or external source of sound. If this phantom sound perception is present over a period of at least six months, it is considered as chronic [Mazurek et al., 2010]. About 10-15% of the global population suffers from tinnitus, whereas in 1-2% it represents a severe burden [Langguth et al., 2013; Heller, 2003; Erlandsson and Dauman, 2013] with comorbidities such as depression, anxiety disorder, sleep disorder or reduced quality of life [Croenlein et al., 2016; Nondahl et al., 2007; Weidt et al., 2016; Trevis et al., 2016].

Currently there is no treatment option for tinnitus available. A major challenge towards an identification of a treatment is related to heterogeneity in tinnitus phenotypes [Hesse, 2016; Kleinjung and Langguth, 2020; Cederroth et al., 2019; Zenner et al., 2017]. Up to now, cognitive behavioral therapy represents the treatment option with the best available evidence for tinnitus [Landry et al., 2020; Cima et al., 2012; Li et al., 2019; Fuller et al., 2020].

In the majority of cases, tinnitus develops as a consequence of cochlear damages subsequent to noise trauma or hearing loss (HL) [Langguth et al., 2013]. Typically, the perceived tinnitus pitch corresponds to the frequency range of maximum HL [Basile et al., 2013; Roberts et al., 2008; Norena et al., 2002; Schecklmann et al., 2012]. Theories about the generation of tinnitus commonly suggest that the reduced or missing auditory input triggers maladaptive alterations along the auditory pathway and the central auditory system, which may lead to the sensation of a phantom sound in the frequencies of the peripheral HL [Eggermont, 2007; Eggermont and Roberts, 2012; Eggermont and Tass, 2015; Adjamian et al., 2009].

On a macroscopic level tinnitus was associated with anomalous oscillatory brain activity patterns such as enhanced activity in the delta and gamma frequency range alongside with reduced alpha activity over temporal regions [Weisz et al., 2005, 2007b]. As observed in several neurophysiological investigations, this delta increase and alpha decrease appears to be closely linked to tinnitus perception as well as tinnitus distress [Weisz et al., 2005; Schlee et al., 2014; Adjamian et al., 2012; Moazami-Goudarzi et al., 2010; Balkenhol et al., 2013]. Due to relations with tinnitus loudness as defined via tinnitus pitch matching [Balkenhol et al., 2013], subjective tinnitus loudness [van der Loo et al., 2009; De Ridder et al., 2015a] or tinnitus-specific increased activity in the auditory cortex [Ashton et al., 2007; Vanneste et al., 2011], high gamma activity was proposed to represent the oscillatory signature of tinnitus perception per se [Weisz et al., 2007b]. These tinnitus-specific spontaneous brain activity patterns were subsumed under the framework of the thalamo-cortical dysrhythmia model (TCD) [Llinás et al., 1999, 2005; De Ridder et al., 2015b], which was further expanded to the “Synchronization-by-Loss-of-Inhibition-Model” (SLIM) [Weisz et al., 2007a].

Conversely, some studies neither observed altered delta and alpha activity in tinnitus [Ashton et al., 2007], any power spectra differences compared to healthy controls [Zobay et al., 2015] nor correlations between electrophysiology and psychoacoustic or psychosocial tinnitus measures [Pierzycki et al., 2016]. In the same vein, further studies report higher alpha activity in tinnitus [Moazami-Goudarzi et al., 2010], a relationship of enhanced alpha and tinnitus intensity [Meyer et al., 2014] or emphasize the relevance of other frequency bands like beta and theta in neural activity related to tinnitus [Meyer et al., 2014; Moazami-Goudarzi et al., 2010; Balkenhol et al., 2013]. Considering these observations, assumptions about abnormal tinnitus-specific respectively tinnitus-related spontaneous brain activity are not so conclusive as presumed initially.

The phenomenon of short-term tinnitus suppression following acoustic stimulation was first studied almost 50 years ago [Feldmann, 1971,1983]. This phenomenon was defined as “residual inhibition” (RI) and can be observed in 60-80% of tinnitus sufferers, whereby depth and duration of suppression patterns vary among individuals [Roberts et al., 2006; Roberts, 2007; Vernon and Meikle, 2003]. Since that time several experiments already examined the impact of various auditory stimulation techniques on RI. These vary from simple white noise (WN) or pure tones, to the application of specific filters or modulation rates, up to the combination of both modulation techniques applied to WN [Henry et al., 2013; Fournier et al., 2018; Roberts et al., 2008, 2006; Tyler et al., 2014; Neff et al., 2017, 2019a; Reavis et al., 2012; Schoisswohl et al., 2019]. It has been suggested that stimulation intensity, duration, specific modulations as well as stimuli including the individual tinnitus frequency (ITF) facilitate short-term acoustic tinnitus suppression.

Another approach to reduce subjective tinnitus loudness for a longer period of time is provided via long-term stimulation with notch filtered music (individual tinnitus pitch is removed from the signal), referred to as “tailor-made notched music training” (TMNMT). The supposed underlying physiological effect behind TMNMT takes place through an inhibition of frequencies within the notch filter called lateral inhibition. By means of long term applications, maladaptive pathological reorganization of the auditory cortex in tinnitus may be reversed [Pantev et al., 2012; Okamoto et al., 2010].

Nevertheless, little is known about the basic neurophysiological processes behind RI [Roberts, 2007]. Reduced firing rates of neurons in the central auditory pathway are theorized to play a key role in RI [Galazyuk et al., 2017, 2019], which covers subcortical structures of the auditory system. There is a paucity in experimental studies examining oscillatory brain activity after acoustic stimulation or rather during RI. With the help of neuromagnetic measures in one tinnitus subject Kristeva-Feige et al. [1995] observed an increase in low frequency (2-8 Hz) spectral power during RI. Contrary to this observation, single-subject intracranial recordings showed a reduction of low frequency (delta: 1-4 Hz; theta: 4-8 Hz) activity in the auditory cortex during RI. These tinnitus-related low frequency oscillations also interacted with alpha (8-12 Hz), beta (20-28 Hz) and gamma (>30 Hz) activity [Sedley et al., 2015]. Beyond that, tinnitus intensity during RI was identified to be connected to delta (1.5-4 Hz), theta (4-8 Hz) and gamma (30-150 Hz) oscillatory activity in the auditory cortex by the use of single patient measurements of neuromagnetic brain activity. The relevance of auditory gamma band activity for RI respectively tinnitus perception could be further corroborated by means of an inverse correlation with tinnitus intensity exclusively in tinnitus subjects experiencing residual excitation [Sedley et al., 2012]. Kahlbrock and Weisz [2008] evaluated neuromagnetic activity in 10 tinnitus patients experiencing RI, defined as 50% of tinnitus loudness reduction for 30 seconds after stimulation offset. A reduction of delta (1.3-4 Hz) activity in temporal areas was observed during RI, whereas the gamma band (low: 30.5-49 Hz; high: 50.3-70.2 Hz) was not affected. The authors conclude that during a short-term reduction of tinnitus intensity, tinnitus-related abnormal oscillatory activities are temporary reversed resulting in a restored balance of neural inhibitory and excitatory processes. A recent study from King et al. [2020] investigated ongoing electrophysiological brain activity of 30 tinnitus subjects following broad band noise stimulation. 17 participants were able to experience RI, whereby a comparison of RI with a control auditory stimulation condition without the ability to induce RI revealed differences with respect to ongoing brain activity. In detail, the authors report higher power in the alpha and gamma frequency bands over the course of RI compared to the control condition.

To the best of our knowledge, the above mentioned five studies represent the only attempts to investigate resting state oscillatory brain activity in the context of RI. The fact that available findings are inconsistent and that merely two experiments - one utilizing magnetoencephalography (MEG) and one Electroencephalography (EEG) - analyzed spontaneous brain activity during RI on a group level indicates an urgent need for respective research whether it is by means of MEG or EEG. Besides single subject analysis, group level analysis represent a basic pillar in science in order to make more general statements about the investigated population e.g., ongoing brain activity associated with RI.

Previous research utilizing neurophysiological measurements, used only one type of non-personalized sound and did not compare participants with and without RI. In the course of this study we are employing an extended set of modified and personalized noise stimuli targeting putatively differential neural mechanisms (i.e., RI and lateral inhibition). Thus the main purpose of this EEG experiment was to examine oscillatory brain activity changes during RI (pre vs. post) following a stimulation with different types of noise. Moreover we aimed to investigate, whether these changes are related to subjective tinnitus loudness ratings. Since RI is a phenomenon which cannot be induced in all people with tinnitus, differences in spontaneous brain activity between people who reported RI and those who didn’t were analyzed (responders vs. non-responders).

Apart from the efficacy of each used stimulus type in short-term tinnitus suppression on a group level, we hypothesize that filtered noise would result in stronger suppression patterns compared to unfiltered noise. In detail, bandstop-filtered noise is assumed to produce the strongest effect via a potentially suppression of neurons reacting to frequencies within the filter range as already shown in long-term applications via TMNMT [Pantev et al., 2012; Okamoto et al., 2010].

Due to the lack of past research in this field, we have no direct stimulus-specific a priori hypothesis about the types of changes from pre to post auditory stimulation in on-going brain activity. However, we assume that potential changes in spontaneous brain activity can be associated with subjective tinnitus loudness ratings after stimulation. In accordance to Kahlbrock and Weisz [2008] we expect a decrease in delta and gamma activity as well as an increase in alpha activity from pre to post auditory stimulation in tinnitus cases experiencing RI (responders). Further we anticipate spectral power differences in the respective frequency bands between acoustic stimulation responders and non-responders. In order to link these differences to auditory cortical activation, source localization of the EEG data was performed.

## 2 Methods

### 2.1 Participants

In the course of this study, N = 45 (14 female) patients with chronic subjective tinnitus (> 6 months tinnitus duration) were recruited from the Interdisciplinary Tinnitus Centre Regensburg, Germany. For participation, patients had to fulfill the following primary inclusion criteria: age between 18 and 75 years; absence of other causes for tinnitus e.g., Meniere’s disease, otosclerosis or acoustic neurinoma; no infection of the oropharynx; no present somatic, neurological or psychiatric disorder; no intake of psychoactive medication (e.g., antidepressants or anticonvulsant drugs), respectively substance or alcohol abuse at least 12 weeks before the start of the experiment; no hypersensitivity to sound; no tinnitus frequency < 1 kHz; no concurrent participation in other tinnitus-related studies or start of any other tinnitus-related treatment in the last three months prior study start.

Ethical clearance with respect to methodological approach and design was sought from the ethics committee of the University of Regensburg, Germany before commencing the experiment (ethical approval number: 17-819-101). For a detailed descriptive overview and clinical characteristics of the sample see table 1. All participants received detailed information about objective, methods, duration and potential side effects of the study. Every participant gave written informed consent before the start of the study and received an appropriate expense allowance after completion of the experiment.

**Table 1:** Sample characteristics. M = mean; SD = standard deviation; Md = median; Min = minimum; Max = maximum; LDL = Loudness Discomfort Level (missings in LDL are due to values over 90 dB); TQ = Tinnitus Questionnaire; THI = Tinnitus Handicap Inventory; VAS = Visual Analog Scale; GUF = Questionnaire on Hypersensitivity to Sound.

| N (female) | 45 (14) |  |  |  |
| --- | --- | --- | --- | --- |
| Tinnitus side (left/ right/ bilateral) | (5/ 8/ 32) |  |  |  |
| Tinnitus loudness fluctuation (yes/ no) | (24/ 21) |  |  |  |
| Tinnitus maskability (yes/ no/ don't know) | (31/ 5/ 9) |  |  |  |
| Musician (yes/ no) | (4/ 41) |  |  |  |
|  | M ± SD | Md | Min | Max |
| Age (years) | 52.29 ± 11.81 | 55.00 | 23.00 | 69.00 |
| Tinnitus duration (months) | 111.04 ± 72.90 | 96.00 | 18.00 | 280.00 |
| Tinnitus frequency (Hz) | 6251.09 ± 2811.38 | 5887.00 | 1020.00 | 15524.00 |
| Tinnitus loudness (dB SPL) | 51.38 ± 16.05 | 50.00 | 27.00 | 85.00 |
| Hearing loss left (dB) | 17.26 ± 13.61 | 14.69 | -5.72 | 55.00 |
| Hearing loss right (dB) | 17.48 ± 11.52 | 17.43 | -8.71 | 45.87 |
| LDL left (dB) (25 missing values) | 86.25 ± 3.21 | 85.50 | 81.00 | 90.00 |
| LDL right (dB) (28 missing values) | 85.06 ± 3.96 | 87.00 | 78.00 | 90.00 |
| Minimum masking level (dB) | 63.82 ± 14.60 | 60.00 | 37.00 | 90.00 |
| Sensation Level (dB) | 47.58 ± 17.49 | 45.00 | 21.00 | 86.00 |
| TQ total score (0-84) | 40.73 ± 15.70 | 40.00 | 17.00 | 71.00 |
| THI total score (0-100) | 35.91 ± 21.38 | 34.00 | 4.00 | 80.00 |
| VAS awareness (%) | 64.62 ± 29.62 | 70.00 | 8.00 | 100.00 |
| VAS loudness (%) | 61.11 ± 24.19 | 65.00 | 15.00 | 100.00 |
| VAS bothersome (%) | 38.20 ± 29.29 | 30.00 | 0 | 100.00 |
| GUF total score (0-45) | 10.73 ± 6.45 | 10.00 | 0 | 23.00 |

### 2.2 Psychometry

Prior to the start of the experiment, participants were requested to answer a set of questionnaires compiled of German versions of the Tinnitus Handicap Inventory (THI) [Newman et al., 1994; Kleinjung et al., 2007], the Tinnitus Questionnaire (TQ) [Goebel and Hiller, 1994; Hallam et al., 1988], the Tinnitus Sample Case History Questionnaire (TSCHQ) [Langguth et al., 2007], visual analog scales (VAS, %) for tinnitus awareness, loudness and bothersome, as well as the Questionnaire on Hypersensitivity to Sound (GUF) [Blaesing et al., 2010] (participants with a score of > 23, which constitutes a very severe impairment, were excluded from our analysis). The survey was performed with SoSci Survey [Leiner, 2016].

### 2.3 Audiometry

Participants hearing thresholds were examined with the toolbox MultiThreshold (University of Essex, United Kingdom) using the implemented paradigm absolute threshold (absThreshold) in Matlab (Matlab R2017a; Mathworks, USA). This paradigm is an implementation of the two-alternatives forced-choice threshold estimation algorithm by Green [1993]. Sine tones (0.5 seconds) were used to test participants hearing level for frequencies from 250 up to 8000 Hz on an octave scale for each ear separately. Starting loudness level was 30 dB SPL, which was increased by 10 dB steps until the participants were able to perceive the sound. The loudness level was raised by 2 dB steps between trials. ER-2 Insert Earphones (Etymotic Research Inc., USA) together with an external soundcard (RME Fireface UCX; Audio AG, Germany) were used for hearing assessment, subsequent matching of the ITF, definition of the sensation level (SL), minimum masking level (MML) (compare section 2.4) as well as the proper auditory stimulation.

### 2.4 Tinnitometry

Individual tinnitus pitch matching was carried out using a Method of Adjustment approach modified from Henry et al. [2013] and Roberts et al. [2008] and implemented in a custom software tool (MAX 7; Cycling’74, USA). A custom-built hardware controller was used comprising a Teensy 3.2 USB-based micro-controller (PJRC, USA) and industrial-grade rotating knobs, switches and motor faders. Detailed information about the used tinnitus matching procedure is described in Neff et al. [2019b]. The starting frequency was defined as one frequency group below the frequency with the highest HL and a start loudness of 10 dB above the particular hearing threshold. Participants tried to match their tinnitus four times as good as possible and rated the accordance of the matched sound with their perceived tinnitus on a 1-10 scale (1 = no accordance; 10 = perfect accordance) after each attempt. The tinnitus matching trial with the highest rating was subsequently defined as the participants ITF. If participants rated different matching attempts similarly, the frequency closest to the mean frequency of the four attempts was chosen. The ITF was then used for the evaluation of further audiometric parameters. Similarly, the MML was defined by increasing the loudness of WN to the point of complete tinnitus masking. Assessment of the loudness discomfort level (LDL) of participants ITF was executed with the discomfort paradigm of the MultiThreshold toolbox with Sennheiser HDA 2000 headphones (Sennheiser, Germany).

### 2.5 Acoustic stimulation

Three different types of noise stimuli with a duration of three minutes each were created in Matlab (Matlab R2017a; Mathworks, USA) with an intensity of 65 dB SL (defined as the loudness level of participants first-time tinnitus pitch perception; maximum loudness of 85 dB SPL) for acoustic stimulation. For this purpose a genuine WN was used to produce individualized noise stimuli through the implementation of bandpass (IBP) and bandstop (IBS) filters with one octave width around the ITF [Pantev et al., 2012]. Each stimuli was composed of a 1000 ms linear fade-in and fade-out phase and underwent a root-mean-square correction to balance levels between stimuli. Diotic acoustic stimulation was performed at a maximum loudness of 85 dB SPL and each stimuli was presented only once. The presentation sequence of the stimuli was randomized.

Before and after the presentation of each stimuli (3 minutes), participants were requested to sit quietly, focus on a white fixation cross on a black screen and avoid extensive eye-blinks and movements while their brain activity was recorded via EEG for three minutes respectively (compare section 2.7).

After the presentation of each noise stimulus, patients had to rate the loudness of their tinnitus at seven different time points (0sec, 30sec, 60sec, 90sec, 120sec, 150sec and 180sec after stimulation offset) on a customized keyboard strip (X-Key-Stick-16-USB, XK-0981-UCK16-R; P.I. Engineering, USA) with a numeric rating scale from 0%to 110%, whereas 100% signified no tinnitus loudness changes, 0% a total absence of tinnitus and 110% an tinnitus loudness increase by 10 %. For an illustration of the acoustic stimulation procedure please see figure 1. The whole experimental stimulation procedure was implemented with the Psychophysics Toolbox Version 3 [Brainard, 1997; Kleiner et al., 2007] in Matlab (Matlab R2017a; Mathworks, USA) and double-blinded. At the end of the experiment, the three stimuli were again presented in a randomized order for 10 seconds each and participants were requested to rate the valence and the arousal of each stimuli via pictorial manikin scales [Bradley and Lang, 1994] on a 9-point Likert Scale, whereas the value 0 indicated a neutral stimulus evaluation (Valence: −4 unpleasant, 4 pleasant; Arousal: −4 relaxing, 4 upsetting).

**Figure 1:**
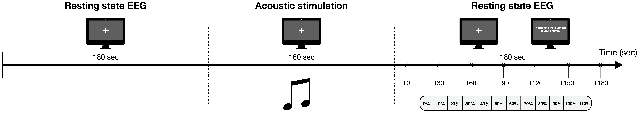
Acoustic stimulation procedure. Prior and post of acoustic stimulation (3 minutes), participants resting state brain activity was recorded via EEG (3 minutes). Participants were instructed accordingly and requested to focus on a white fixation cross on a black screen during the whole experiment. Following acoustic stimulation, participants were requested to rate the current loudness of their tinnitus (“Please rate the loudness of your tinnitus.”) at seven points in time (0, 30, 60, 90, 120, 150 and 180 seconds towards stimulation offset) on a numeric rating scale from 0% to 110% (0% - total absence of tinnitus; 100% - no tinnitus loudness changes; 110% −10% tinnitus loudness increase). This acoustic stimulation procedure was repeated for each of the three used types of noise stimuli (white noise, individualized bandpass filtered white noise, individualized bandstop filtered white noise).

### 2.6 Behavioral Analysis

Behavioral data was analyzed with the statistic software R (R version 3.4.2; R Foundation for Statistical Computing, Austria) and the packages “psych”, “emmeans”, “sjstats” and “lme4”. Linear mixed effect models were used to analyze tinnitus loudness ratings and stimuli evaluation (valence, arousal) separately. The following predictors were tested for the model fitting procedure of tinnitus loudness ratings: condition (stimuli, compare section 2.5), time (0sec, 30sec, 60sec, 90sec, 120sec, 150sec, 180sec towards stimulation offset), tinnitus bilaterality (yes/no), sex (male/female), tinnitus duration and stimuli position in the auditory stimulation sequence. The predictors condition, gender and tinnitus duration were tested for the model fitting procedure of stimuli evaluation data.

Other potential predictors such as tinnitus loudness (dB), MML, SL or HL were not included in the model fitting procedure, since they were experimentally controlled e.g., by the creation of tailored stimuli. Participant (id) was considered as a random effect in all model fitting procedures. In order to identify the model with the best fit for the data, the step function of the lme4 package was deployed. Thereby, a backward elimination of non significant predictors as well as a forward addition of significant predictors is conducted by comparing the models with Likelihood Ratio Tests [Harrison et al., 2018]. Marginal (variance of the predictors) and conditional (variance of predictor and random effect) *R*^2^ were computed to provide the amount of the explained variance of the respective model [Nakagawa et al., 2017]. For each final model, fixed effects were examined via Expected Mean Square Approach. Potential differences in tinnitus loudness and stimuli evaluation within predictors were analyzed with post-hoc Tukey-tests. Analysis of descriptive differences between HL and LDL between the left and right ear were tested by the means of two-sample t-tests. Normal distribution (Shapiro-Wilk-Test) and homoscedasticity (F-test) were examined and if violated, non-parametric testing with independent sample Mann-Whitney U-tests were conducted. To evaluate effect size of significant differences, Cohen’s d was calculated. The level of statistical significance was set to p ≤ .05 for all analyses.

### 2.7 Electrophysiological data acquisition and analysis

#### 2.7.1 EEG recording

EEG data was recorded with a BrainAmp DC system, EasyCap electrode cap with 64 electrodes, and Brain Vision Recorder 1.20 software (Brain Products GmbH, Germany). The sampling rate was 500 Hz and electrodes were referenced to FCz during recording. Impedances were kept below 10kΩ.

#### 2.7.2 Preprocessing

Raw EEG data was preprocessed with a custom-built semi-automatic pipeline using the Fieldtrip toolbox [Oostenveld et al., 2011] in Matlab (Matlab R2017a; Mathworks, USA). EEG data was filtered between 0.5 Hz and 45 Hz with a 4th order Butterworth bandpass filter.

Hereafter, an independent component analysis (ICA, fastICA http://research.ics.aalto.fi/ica/fastica/index.shtml) was used to identify and remove components with horizontal and vertical eye movement. Noisy or aberrant channels were interpolated using weighted neighbors. Neighboring channels were defined via a triangulation of 2D sensor position projection and channels identified for interpolation were replaced with the mean of neighboring sensors. In a next step, average referencing was performed and the recording reference electrode FCz was added as a data channel. In order to control for noisy channels introduced by the rating procedure of the post stimulation conditions, posterior (Iz, TP9, TP10) as well as frontal channels (FPz, FP1, FP2, AF3, AF4, AF7, AF8) were discarded from subsequent analyses steps. Data was then segmented into 2 seconds segments. All segments during which participants rated the loudness of their tinnitus were rejected. Additionally, one segment before and after the rating was excluded as well. Segments with remaining artifacts were rejected with combined automatic identification via a z-score (*μ*V) threshold of −2/ +2 and visual inspection in a final step. Average number of valid segments was different (U = 1970.50, p = .001) between pre (M = 78.93, SD = 6.48) and post (M = 60.37, SD = 6.19) acoustic stimulation.

#### 2.7.3 EEG analysis

##### Power analysis - whole group

Frequency power spectra of pre and post auditory stimulation datasets per subject and condition(compare 2.5) were calculated using multitaper frequency transformation (mtmfft) and a hanning window with a spectral smoothing of 1 Hz. Next, grand averages were created for pre and post stimulation datasets per condition by computing power spectra averages across all valid segments and all subjects.

Potential changes in EEG power spectra were analyzed with a 2 x 3 repeated measurement ANOVA and the within subject factors time (pre, post) and condition (WN, IBP, IBS), which was implemented in Fieldtrip. The main effects for time and condition were tested with paired two-sided t-tests via non-parametric cluster-based permutation tests with 10.000 iterations. In order to test for an interaction effect of time and condition, a dependent samples multivariate ANOVA was conducted using a non-parametric cluster-based permutation test with 10.000 iterations as well. We were primary interested in an interaction effect of time and condition. In case of a significant time x condition interaction, effects were followed up using post-hoc contrasts. Pre vs. post contrast per condition were analyzed with dependent samples t-tests, whereas potential differences in stimuli-induced power spectra changes from pre to post stimulation as well as post stimulation differences (inter-stimulus contrasts), were contrasted via independent samples t-tests using non-parametric cluster-based permutation test as described above.

Additionally, Pearson correlations between post stimulation power spectra and prepost power spectra differences with averaged tinnitus loudness ratings (over all 7 time points) as well as directly after stimulation offset (T0) were computed via cluster-based permutation tests. Significance level was set to p ≤ .05 for all EEG analyses and p <0.1 was defined as a statistical trend. Significant clusters were defined as a minimum of two significant neighboring channels for all analysis. For the purpose of interpretation, EEG frequency bands were defined as follows: delta 1-4 Hz, theta 5-7 Hz, alpha 8-12 Hz, beta 13-29 Hz, gamma 30-45 Hz.

##### Power analysis - responder

Furthermore, we compared frequency power spectra of participants who exhibited RI with those who did not experience RI after auditory stimulation. For this purpose RI was defined as ≤ 50% of tinnitus loudness directly after stimulation offset resulting in a subset of n = 12 further indicated as responders. Within this subgroup of responders, n = 5 participants each, responded to a stimulation with WN or IBP, whereas only n = 2 participants reported RI after a stimulation with IBS. A second subgroup of participants without RI (non-responders) were matched to responders according to the following criteria: gender; mean HL; age and absence of RI (tinnitus loudness of ≥ 100% after stimulation offset) in the same stimulus type as matched patient exhibited RI in responders group. Sample characteristics for both subgroups can be seen from table 2. Associations of categorical variables with stimulation response (responder or non-responder) were analyzed with χ^2^-tests or Fisher’s exact tests if cell frequencies were below 5. Differences in numerical variables between the two subgroups were analyzed by two-sample t-tests. In case of violated statistical assumptions, Mann-Whitney U-tests were performed. Significance levels were set to p ≤ .05 and a statistical trend was defined as p <0.1.

**Table 2:**
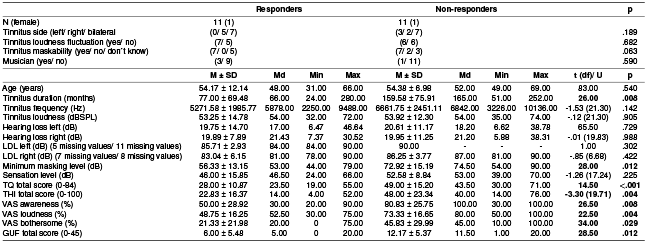
Sample characteristics - responders vs. non-responders. M = mean; SD = standard deviation; Md = median; Min = minimum; Max = maximum; df = degrees of freedom; LDL = Loudness Discomfort Level (missings in LDL are due to values over 90 dB); TQ = Tinnitus Questionnaire; THI = Tinnitus Handicap Inventory; VAS = Visual Analog Scale; GUF = Questionnaire on Hypersensitivity to Sound.

Power spectra for pre and post auditory stimulation EEG datasets were averaged over all subjects within the respective subgroup (responders and non-responders). Analysis were conducted using normalized EEG datasets by dividing power spectra for each single frequency through the mean power of the entire frequency spectrum.

Illustrated power spectra per frequency were transformed according to 10 * log10(x). EEG power spectra were analyzed with a 2 x 2 repeated measures ANOVA and the factors time (pre, post) and group (responders, non-responders). The main effects for time and group were evaluated with dependent sample respectively independent sample t-tests according to the same approach as already described in the power analysis section for the whole group. Likewise, a potential interaction effect of time and group was analyzed with an independent samples t-test.

In the case of a significant interaction effect, post-hoc dependent samples t-tests for pre vs. post within subgroup contrast and independent samples t-tests for between subgroup contrast (responders vs. non-responders) separated for pre and post stimulation measurements are conducted. Regardless of an observed interaction effect, an exploratory contrast of post stimulation power spectra differences between responders and non-responders is performed. Equal to the whole group analysis, Pearson correlations were calculated with cluster-based permutation tests for post stimulation power spectra and pre-post power spectra differences with averaged tinnitus loudness ratings or rather directly after stimulation offset (T0). Additionally, a correlation of post stimulation power spectra and pre-post power spectra differences with tinnitus loudness rated via VAS (%) was computed.

In order to explore differences in cortical alpha variability between responders and non-responders a coefficient of variance was calculated by dividing the standard deviation of the alpha frequency power (8-12 Hz) by its mean power.

##### Source space analysis

Source localization of frequency data was performed using a standard boundary element headmodel [Oostenveld et al., 2003] and the dynamic imaging of coherent sources algorithm optimized for EEG frequency data (Dynamical Imaging of Coherent Sources, [Groß et al., 2001]). Inter-subgroup source contrasts (responders vs. non-responders; responders vs. non-responders post stimulation) of peak frequencies received from sensor-level cluster analysis (maximum value) were analyzed via non-parametric cluster-based permutation tests with 10.000 iterations using normalized EEG datasets. Normalization procedure was identical to the sensor level analysis.

## 3 Results

### 3.1 Sample characteristics

Table 1 summarizes the descriptive statistics and tinnitus-related questionnaire scores of the present sample. In the majority of participants, tinnitus was perceived bilaterally (n = 32) and featured loudness fluctuations (n = 24). The possibility to mask their perceived tinnitus was reported by n = 31 participants. Moreover, n = 4 participants claimed to be musicians and the average duration of tinnitus perception was 111.04 months (SD = 72.90).

Stimulation with either WN, IBP and IBS resulted in n = 12 responders, who showed RI with at least one stimulus type.

A weak association of stimulation response (responders or non-responders) and tinnitus maskability (yes, no, don’t know) was found with the group of responders exhibiting no participant who reported an absence of tinnitus maskability (cf. table 2). Statistical testing for differences between the subgroups of responders and non-responders revealed differences in terms of tinnitus duration, MML and questionnaire data with the group of responders showing shorter tinnitus duration (U = 26.00, p = .008, d = 1.135), lower MML (U = 28.00, p = .012. d = 1.168) as well as lower sum scores in TQ (U = 14.50, p <.001, d = 1.159), THI (t (19.71) = −3.30, p = .004, d = 1.249) and GUF (U = 28.50, p = .012, d = 1.137). Likewise, responders reported lower values in subjective measurements of tinnitus awareness (U = 26.50, p = .008, d = 1.126), loudness (U = 22.50, p = .004, d = 1.494) and bothersome (U = 34.00, p =.029, d = .931) as indicated by VAS (in %). Detailed sample characteristics and statistical comparisons for the two subgroups are shown in table 2.

### 3.2 Audiometry and Tinnitometry

Results from audiometric assessment and tinnitus matching are outlined in table 1 as well as illustrated in figure S1. The investigated sample featured a mean tinnitus frequency of 6251.09 Hz (SD = 2811.38), whereas the average tinnitus loudness was 51.38 dB SPL (SD = 16.05). Initial perception of the individual tinnitus pitch (SL) appeared at a mean volume level of 47.58 dB (SD = 17.49). Mann-Whitney U-tests found no differences with respect to HL (U = 941.50, p = .569) and LDL (U = 199.50, p = .361) between the left and the right ear.

### 3.3 Acoustic Stimulation

Table S1 lists the descriptive statistics for tinnitus loudness ratings for each stimuli on average as well as time point T0. Tinnitus suppression time curves, including all seven time points, are illustrated in figure 2 for each stimuli.

**Figure 2:**
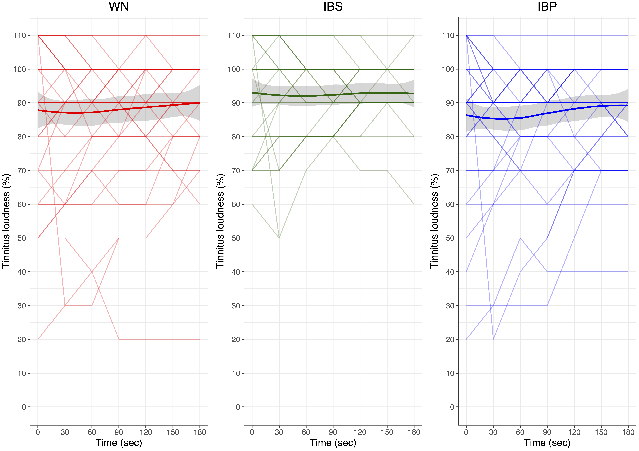
Tinnitus loudness time curve per condition. WN = white noise; IBP = individualized bandpass filtered white noise; IBS = individualized bandstop filtered white noise. Tinnitus loudness ratings are illustrated on a single participant level for all rating timepoints separated for each stimuli. Thick lines show the mean tinnitus loudness (%) per stimulus, standard deviations are illustrated as grey ribbons.

Model fitting procedure of behavioural data was able to identify the following model with the best fit for the data: *response ~ condition* + (1 | *id).* Table S2 lists detailed results of the model fitting proceeding. A significant effect of condition was observed (cf. table S3). Suceeding post-hoc contrasts found differences between stimulus WN vs. IBS, as well as IBP vs. IBS (cf. table 3). A potential confounding caused by the position of the stimuli in the acoustic stimulation sequence could be excluded, since position did not appear as a significant predictor in the final model.

**Table 3:** Post-hoc tukey contrasts for condition. WN = white noise; IBP = individualized bandpass filtered white noise; IBS = individualized bandstop filtered white noise; degrees of freedom = 902.00; standard error = .87.

| Contrast | Estimate | t | p | d |
| --- | --- | --- | --- | --- |
| **Total sample** |  |  |  |  |
| WN - IBP | 1.05 | 1.20 | .451 | .057 |
| **WN - IBS** | **-4.32** | **-4.96** | **<.001** | **.251** |
| **IBP - IBS** | **5.37** | **-6.17** | **<.001** | **.328** |

### 3.4 Stimulus evaluation

Stimulus evaluation outcomes in terms of valence and arousal can be seen from table S4 and figure S2. Model *response ~ condition* + (1 | id) was identified to have the best fit for the valence data with condition as a significant fixed effect (cf. tables S5 and S6). Post-hoc tests were able to reveal differences for valence evaluations of stimuli WN vs. IBS and also IBP vs. IBS as can be seen from table S7. Subsequent model was identified by our model fitting approach for arousal data: *response ~ condition* + *gender* + (1 | *id)* (cf. table S5). Fixed effect testing revealed significant effects for condition and gender (cf. table S6). Post-hoc analysis showed differences between stimuli IBP and IBS as well as male and female participants (cf. table S7).

### 3.5 Electrophysiology

Results of whole sample EEG power spectra analysis are outlined in table 4. A significant main effect of time was observed, indicating higher spectral power for 1-7 Hz and 26-45 Hz plus lower spectral power for 7-28 Hz after auditory stimulation. Further, a significant interaction of condition and time was found in the frequency spectra 1-7 Hz and 36-45 Hz. Succeeding post-hoc contrasts revealed higher power in lower frequencies towards stimulation across all stimuli (WN: 1-7 Hz; IBP: 1-6 Hz; IBS: 1-6 Hz) as well as higher gamma activity after a stimulation with IBP (32-45 Hz) and IBS (37-45 Hz). A power decrease following IBS stimulation was found for the frequency cluster 11-19 Hz. In addition, statistical trends towards power reductions in the frequency clusters 10-12 Hz and 14-19 Hz were observed for pre-post comparisons of stimulus WN. Differences between the applied types of stimuli with respect to pre-post power spectra changes or post stimulation power spectra were not detected.

**Table 4:** Electrophysiology - results of cluster-based permutation test for the total sample analysis. WN = white noise; IBP = individualized bandpass filtered white noise; IBS = individualized bandstop filtered white noise; df = degrees of freedom; Max = maximum. Positive clusters indicate increased power spectra whereas negative clusters indicate decreased power spectra from pre to post stimulation, in the respective frequency ranges. Peak frequency (Hz) and peak electrode represent the particular frequency and electrode featuring the maximum value obtained from cluster statistics.

|  | Frequency (Hz) | Cluster statistic (df) | p | Peak frequency (Hz) | Peak electrode | Max. statistic |
| --- | --- | --- | --- | --- | --- | --- |
| **Time** |  |  |  |  |  |  |
| Positive cluster | 1-7 | t(134) = 1047.88 | <.001 | 4 | PO8 | 7.68 |
| Positive cluster | 26-45 | t(134) = 893.13 | <.001 | 41 | POz | 4.89 |
| Negative cluster | 7-28 | t(134) = -1150.64 | <.001 | 12 | T8 | -5.33 |
| **Condition x Time** |  |  |  |  |  |  |
| Positive cluster | 1-7 | F(5,40) = 3437.77 | .002 | 4 | PO8 | 51.28 |
| Positive cluster | 36-45 | F(5,40) = 2783.52 | .002 | 42 | F6 | 34.09 |
| **Post-hoc - pre vs. post stimulation per stimulus** |  |  |  |  |  |  |
| **Positive cluster** |  |  |  |  |  |  |
| WN | 1-7 | t(44) = 482.28 | .006 | 5 | O1 | 4.81 |
| IBP | 1-6 | t(44) = 696.17 | .002 | 3 | O1 | 5.90 |
| IBP | 32-45 | t(44) = 460.98 | .007 | 41 | F3 | 4.44 |
| IBS | 1-6 | t(44) = 398.13 | .006 | 3 | O2 | 4.20 |
| IBS | 37-45 | t(44) = 199.09 | .026 | 45 | P2 | 3.54 |
| **Negative cluster** |  |  |  |  |  |  |
| WN | 10-12 | t(44) = -132.92 | .058 | 11 | T8 | -4.24 |
| WN | 14-19 | t(44) = -123.90 | .064 | 19 | C3 | -3.95 |
| IBS | 11-19 | t(44) = -242.31 | .016 | 13 | T8 | -4.20 |

Electrodes within frequency clusters as outlined in table 4 can be found in the supplemental material in table S8 grouped by brain areas.

No correlations were found on the cluster level for post stimulation EEG power or pre-post power spectra changes with averaged tinnitus loudness ratings or rather tinnitus loudness ratings immediately after stimulation end (T0) for any of the used stimuli.

Table 5 provides the results obtained from the responder EEG power spectra analysis (compare section 2.7.3). A significant main effect of time was observed, indicating a power reduction from pre to post stimulation in the frequency cluster 6-32 Hz for responders as well as non-responders. Likewise, a significant effect of group demonstrates lower power in higher frequency ranges (22-45 Hz; t(max) = −4.06, over electrode P5 at 31 Hz; cf. figure 3 A and B) as well as a statistical trend towards higher power in the alpha frequency range (7-12 Hz; t(max) = 4.35, over electrode F4 at 9 Hz; cf. figure 3 A and B) for the subgroup of responders. There was no significant interaction of time and group. Electrodes within frequency cluster presented in table 5 can be found in table S9 in the supplemental material.

**Table 5:** Electrophysiology - results of cluster-based permutation test for the responder analysis. df = degrees of freedom; Max = maximum. Positive clusters indicate increased power spectra, whereas negative clusters indicate decreased power spectra for responders compared to non-responders respectively from pre to post stimulation (effect of time) in the respective frequency ranges. Peak frequency (Hz) and peak electrode represent the particular frequency and electrode featuring the maximum value obtained from cluster statistics.

|  | Frequency (Hz) | Cluster statistic (df) | p | Peak frequency (Hz) | Peak electrode | Max. statistic |
| --- | --- | --- | --- | --- | --- | --- |
| **Time** |  |  |  |  |  |  |
| Negative cluster | 6-32 | t(11) = -1539.00 | <.001 | 18 | TP7 | -6.77 |
| **Group** |  |  |  |  |  |  |
| Positive cluster | 7-12 | t(22) = 246.27 | .082 | 9 | F4 | 4.35 |
| Negative cluster | 22-45 | t(22) = -573.34 | .024 | 31 | P5 | -4.06 |
| **Exploratory post-hoc contrast - responders vs. non-responders post stimulation** |  |  |  |  |  |  |
| Positive cluster | 5-17 | t(22) = 549.39 | .035 | 9 | F4 | 4.94 |

**Figure 3:**
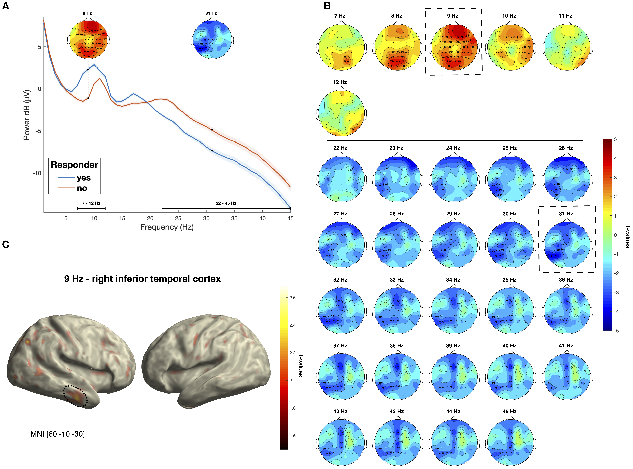
Responders vs. non-responders - contrast of power spectra at the sensor and source level. **A**: Power spectra differences for responders and non-responders for the frequencies 1-45 Hz. Significant positive cluster 5-17 Hz and negative cluster 22-45 Hz as well as the respective peak frequencies (9 Hz and 31 Hz) are highlighted. Grey ribbons represent the standard deviation for each subgroup. **B**: Cluster statistic results (t-values) of power spectra contrasts between responders and non-responders are presented as topographic plots per frequency for a positive cluster of 5-17 HZ and a negative cluster of 22-45 Hz. Significant cluster electrodes are accentuated in bold and labeled per frequency. Peak frequencies of 9 Hz and 31 Hz, representing the maximum values obtained from the cluster statistics, are highlighted with dashed line rectangles. **C**: Source localization of 9 Hz EEG power peaking in the right inferior temporal gyrus (BA 20).

Coefficient of variance calculation exclusively for the alpha frequency band (8-12 Hz) exposed a higher variation in frequency band power for the subgroup of responders (responders: 61.04%; non-responders: 50.03%)

Correlations of EEG power towards stimulation or pre-post power spectra changes on the cluster level with subjective tinnitus ratings for the group of responders showed no significant results for mean tinnitus loudness or tinnitus loudness at T0. Further no correlation with tinnitus loudness rated via VAS (%) was observed.

Subsequent exploratory analysis of post stimulation power spectra differences between responders and non-responders, exhibited increased activity in the frequency cluster 5-17 Hz in the subgroup of responders (t(max) = 4.94, over electrode F4 at 9 Hz; cf. table 5 and figure 4 A and B).

**Figure 4:**
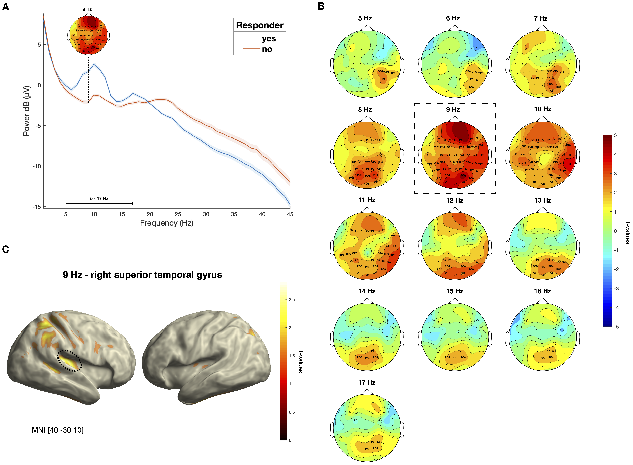
Responders vs. non-responders - post stimulation power spectra contrasts at the sensor and source level. **A:** Power spectra differences for responders and non-responders for the frequencies 1-45 Hz. Significant positive cluster 5-17 Hz with the respective peak frequency of 9 Hz is highlighted. Grey ribbons represent the standard deviation for each subgroup. **B**: Results of cluster statistics (t-values) of power spectra contrasts between responders and non-responders following acoustic stimulation are presented as topographic plots per frequency for a positive cluster comprised of 5-17 Hz. Significant cluster electrodes are accentuated in bold and labeled per frequency. Peak frequency of 9 Hz is highlighted with a dashed line rectangle. **C**: Source localization of 9 Hz EEG power peaking in the right superior temporal gyrus (BA 41).

Projecting peak frequencies of sensor-level power differences of responders and non-responders contrasts in source space exposed differences solely for 9 Hz (t(cluster) = 13.07, p = .004) with maximum differences (t(max) = 2.70) localized in the right inferior temporal gyrus (MNI: 60 −10 −30) shown in figure 3C). However, no difference at the peak frequency 31 Hz could be observed in source space. Source localization of the peak frequency received from sensor-level contrast between responders and non-responders post acoustic stimulation exhibited differences at the frequency of 9 Hz (t(cluster) = 31.95, p = .032) localized in the right superior temporal gyrus (MNI: 40 −30 10) presented in figure 4C.

## 4 Discussion

The main objective of the present study was to investigate the effect of different types of noise stimuli on short-term tinnitus suppression and corresponding electrophysiological brain activity. Moreover, we wanted to elucidate if electrophysiological changes are a function of tinnitus loudness ratings and if differential activation patterns arise from the different stimuli putatively triggering RI or lateral inhibition, respectively. Finally, we aimed at examining potential differences in ongoing brain activity between responders and non-responders. To the best of our knowledge, this presentation of notch- and bandpass-filtered WN sounds is novel in its application in tinnitus research. Similarly, we are the first group which elucidated neurophysiological differences between acoustic stimulation responders and non-responders. In the following, the results of our study are thus critically discussed in the light of current knowledge and with respect to future research outlook.

### 4.1 Behavioral results

The behavioral analysis demonstrate similar suppression patterns as past studies in this field with only a subset of the study population reporting a considerable tinnitus loudness reduction after acoustic stimulation. On a group level all of the used stimuli induced short-term tinnitus suppression. Contrary to our hypothesis IBS appeared to produce the fewest reduction in tinnitus loudness rating, whereas IBP resulted in the strongest suppression pattern.

A potential explanation for this difference might derive from the ability of IBP/ WN in stimulating a broader range of frequencies around the ITF leading to a reduction of neural response gain and tinnitus-related hyperactivity and as a result facilitating short-term tinnitus suppression (cf. Schaette et al. [2010]), whereas suppressing effects of IBS via lateral inhibition might only appear after long-term application.

However, it is also possible that so called feed-forward inhibition is responsible for the superiority of stimuli containing signal in frequency ranges affected by hearing loss (cf. Roberts [2007]; Roberts et al. [2010]).

These explanations remain highly speculative and currently we are not able to provide a suitable explanation for these observed differences. Interestingly, stimulus IBP was evaluated with the lowest tolerability as indicated by the highest arousal and lowest valence ratings. This finding is contrary to one of our previous experiments which reports low arousal and high valence ratings for IBP [Schoisswohl et al., 2019].

Generally, about 50 to 90% of the studied individuals report some level of tinnitus suppression after acoustic stimulation (e.g., [Neff et al., 2017; Schoisswohl et al., 2019; Fournier et al., 2018; Kahlbrock and Weisz, 2008; Sedley et al., 2012]). Given the skewed distribution of RI responses on the group level in previous and this study as well as the need for a reliable threshold for strong tinnitus suppression, we opted to define a reduction in tinnitus of 50% after acoustic stimulation as the threshold for the responder classification akin to [Kahlbrock and Weisz, 2008]. Applying this threshold, we can report an absolute number of 12 responders (with any stimulus type) out of 45 participants (26.67% responder rate) which is comparable to relative numbers reported by Kahlbrock and Weisz [2008] (26% responder rate), but below the quantity of responders reported by King et al. [2020] (56.67% responder rate; the threshold for RI in this study is currently unknown due to publication status).

### 4.2 Electrophysiology

Since only a handful of studies evaluated neural activity during RI, no specific hypotheses were generated about oscillatory changes from pre to post stimulation. In light of past neurophysiological research and the assumptions that tinnitus is accompanied by abnormal delta, alpha and gamma activity [Weisz et al., 2005, 2007a; Adjamian et al., 2012; Moazami-Goudarzi et al., 2010; Balkenhol et al., 2013; van der Loo et al., 2009; Ashton et al., 2007] as well as a putative brief inversion of altered spontaneous brain activity during RI [Kahlbrock and Weisz, 2008], it can be supposed that observed group-level changes in tinnitus loudness (RI) are also reflected in electrophysiological measures. Namely, a reduction in delta and gamma and an increase in alpha power spectra from pre to post stimulation is to be expected given these assumptions.

### 4.3 Whole group analysis

Analysis of whole group pre-post stimulation changes in ongoing brain activity revealed increases in the delta, theta and gamma frequency range as well as decreases in alpha and beta frequency bands. This increase in low frequency activity is in direct contrast to past observations, which report a reduction of delta and theta power spectra during RI in accordance with the current neurophysiological models for tinnitus [Kahlbrock and Weisz, 2008; Sedley et al., 2012, 2015]. In contrast, an earlier study using neuromagnetic measures in a single subject during short-term tinnitus suppression likewise reports an enhancement of low frequency activity [Kristeva-Feige et al., 1995].

Gamma band activity was suggested to represent a spontaneous brain activity pattern related to the actual tinnitus perception [Weisz et al., 2007a], therefore it is assumed that during a potential suppression of tinnitus after acoustic stimulation, activity in the gamma band will be suppressed. The current findings revealed an increase in gamma power after auditory stimulation, similar to findings from Sedley et al. [2015, 2012]; King et al. [2020], who observed an increase in gamma band activity during RI. Consistent with the current literature, we observed a decrease in alpha frequency band power from pre to post stimulation [Kahlbrock and Weisz, 2008; Sedley et al., 2015]. However, a recent study was able to demonstrate an increase in alpha frequency band power during RI in accordance with the given neurophysiological models in tinnitus [King et al., 2020].

No relationship of pre-post power spectra changes, neither with tinnitus loudness ratings averaged over all time points nor directly after stimulation offset was observed in our data. Past neurophysiological research was not able to produce consistent findings in terms of correlations with behavioral measures of tinnitus respectively RI (e.g., intensity, loudness). Besides observed positive correlations of low and high frequency activity [Sedley et al., 2012; Balkenhol et al., 2013; van der Loo et al., 2009] or alpha activity with tinnitus intensity [Sedley et al., 2015; Meyer et al., 2014], the current findings are in accordance with other studies which report an absence of any relationship [Adjamian et al., 2012; Pierzycki et al., 2016; Kahlbrock and Weisz, 2008]. In consideration of missing correlations as well as power spectra changes in conflict with current neurophysiological models for tinnitus, we suggest that the present findings do not indicate oscillatory patterns related to tinnitus loudness suppression, rather constitute a tinnitus-unspecific neurophysiological reaction to an external acoustic stimulus.

Oscillatory activity in the alpha frequency range is supposed to be relevant for inhibitory processes of the brain [Klimesch et al., 2007], thus a sound stimulation exceeding the individual tinnitus loudness level produces excitation and consequently alpha decreases. It has already been shown, that spontaneous activity in the alpha (6-12 HZ) and beta (~20 Hz) frequency bands desynchronize after sound stimulation (for an overview see Weisz et al. [2011]). Likewise, gamma band activity (30-45 Hz; 80-100 Hz), which is associated with cortical activation like attention or perception, was observed to be enhanced after the presentation of sound stimuli [Crone et al., 2001; Joliot et al., 1994] comparable to the present and recent findings [King et al., 2020].

In order to distinguish spontaneous brain activity related to tinnitus suppression from tinnitus-unspecific neurophysiological consequences to a sound stimulation, future research should not only compare acoustic stimulation responders and non-responders (RI vs. absence of RI) but also strive for a comparison with healthy control groups.

### 4.4 Responder analysis

Another objective of this study was to compare acoustic stimulation responders with nonresponders, in order to point out potential differences in regards to ongoing brain activity. To the best of our knowledge this is the first study, which compares oscillatory activity of acoustic stimulation responders and non-responders.

Interestingly, we observed reduced gamma band activity and a trend for enhanced alpha activity (peak frequency of 9 Hz localized in the right inferior temporal gyrus; BA 20) for the group of responders in contrast to non-responders. This result may corroborate the premise that gamma might be related to tinnitus perception [van der Loo et al., 2009; De Ridder et al., 2015a; Ashton et al., 2007; Weisz et al., 2007b]. Given the fact, that responders generally reported their perceived tinnitus loudness level lower than non-responders, the question arises if the perceived tinnitus loudness rated via VAS can be associated with ongoing brain activity e.g., lower tinnitus loudness related to reduced gamma power or enhanced alpha. Yet, a respective correlation analysis failed to show an association.

As already shown by Schlee et al. [2014] tinnitus sufferers exhibited a blunted alpha peak and more importantly reduced alpha variability (8-10 Hz). This finding could be reflected by our data in a similar way as non-responders had a lower alpha peak and lower alpha variability (8-12 Hz). In further support for this argumentation, the data of the former study as well as our present findings show longer tinnitus duration for subjects with reduced alpha power, whereas we assume that these insights from case-control contrasts can be applied to the responder analysis at hand.

The observed reduction in gamma power may be interpreted along similar veins as the findings in alpha power by applying insights from case-control studies. Responders with a less chronified and intense tinnitus in our study are thus comparable to healthy controls in some case-control designs with reported lower gamma power values [Ashton et al., 2007; Vanneste et al., 2011]. In further analogy, our findings of diminished gamma band activity together with a decrease in tinnitus loudness for the subgroup of responders can be linked to observations of past studies, namely a positive correlation of gamma with tinnitus loudness [van der Loo et al., 2009; De Ridder et al., 2015a; Balkenhol et al., 2013].

We theorize that this trend for blunted alpha as well as lower gamma activity may be indicative of a trait as a consequence of tinnitus chronification.

A related observation was made by Neff et al. [2019a] where active listening to tinnitus and consequential increase in tinnitus intensity did not lead to any neural alterations, which fits the reasoning about a trait-like neural representation of chronified tinnitus.

However, it is also possible that this pattern of reduced gamma and enhanced alpha activity represent a genuine neural trait related to acoustic stimulation response more specifically the possibility to induce RI in tinnitus sufferers.

Our exploratory analysis of post acoustic stimulation contrasts revealed higher spectral power in the theta, alpha and beta frequency range with a peak in the alpha band (9 Hz) localized in the right superior temporal gyrus (BA 41) in acoustic stimulation responders.

This increased alpha in auditory fields is in line with our hypothesis of a brief inversion of altered oscillatory power during RI and is consistent with past research examining disparities between tinnitus and healthy controls (compare section 1). Notably, this supports our assumptions about responders and related trait-like neural signatures of tinnitus in that it surmises that only responders can exhibit neural responses which are specific to RI induced by acoustic stimulation.

Finally, a lack of correlations between loudness ratings and ongoing brain activity in the present study does not allow for a conclusive interpretation with regards to tinnitus. Past studies examining correlates of tinnitus suppression and neural activity have been able to demonstrate a relationship of low and high frequency activity with tinnitus intensity Sedley et al. [2015, 2012]. Nevertheless Kahlbrock and Weisz [2008] were not able to demonstrate a correlation of tinnitus suppression and ongoing neural activity in agreement with the present findings.

To further investigate these observed differences it is recommended to optimize future study designs with respect to a parametric analysis of tinnitus duration and RI-related neural activity.

### 4.5 Limitations

Our study has several limitations which might be informative for future research in the specific subfield of acoustic stimulation and general research in tinnitus.

No correlations between neurophysiological changes and changes in behaviorally assessed self-report tinnitus loudness were found in our data. Given the narrow and skewed distribution of the behavioral data and the consequential arbitrary choice of a RI threshold of 50% for the responder group contrast, correlation analysis might neither way be informative with the current data. This negative result is in line with the former study of Kahlbrock and Weisz [2008]. Moreover, full and prolonged RI could only be studied in a small subset of the participants. Finally, heterogeneity of tinnitus loudness suppression curves between participants and the general low reliability and validity of tinnitus self-report data may further contribute to these absent findings.

As in many previous studies, it is challenging to recruit a large enough study sample from the locally available tinnitus population for the extensive experimental procedures. Additionally, tinnitus suppression responses, especially the parameters of RI depth as well as duration, can not be properly assessed in established screening procedures. This selection bias is hard to come by and potentially distorts results. Future studies could thus profit from internet-based prescreening. Beyond that, multi-center studies could help to further increase the validity of results aside from increasing the sample size.

## 5 Conclusions

The main goal of the current study was to unveil the oscillatory signature of RI and see how this relates to established neurophysiological models of tinnitus. In contrast to former studies, we used an extended set of modified noise stimuli targeting putatively differential neural mechanisms (i.e., RI and lateral inhibition). Furthermore, we explicitly investigated responder profiles of RI. Similar to former studies, merely a quarter of tested participants exhibited pronounced RI.

Looking at the oscillatory signature of acoustic stimulation responders and non-responders, results are indicative of decreased gamma and increased alpha power for responders. These findings are in line with both the proposed models of SLIM and TCD, respectively. This observations might be indicative of trait-specific forms of oscillatory signatures in different subsets of the tinnitus population possibly related to acoustic tinnitus suppression. In agreement with a potential transient reversal of tinnitus-specific abnormal ongoing brain activity over the course of tinnitus suppression, alpha power was enhanced in the group of responders after stimulation similarly compared to non-responders. Source localization of the sensor-level differences emphasizes the involvement of auditory cortical systems. Given the lack of correlations between tinnitus loudness and oscillatory power in this study, which was also reported by former studies, results do not allow for a conclusive interpretation with respect to these models.

The identified tinnitus patient profile experiencing RI, which mainly features less tinnitus chronification, could serve as a selection criterion to identify individuals for successful acoustic tinnitus suppression and putatively for acoustic treatments (e.g., treatment start in early stages of chronification).

Further research examining oscillatory activity during RI should strive for a healthy control group as well as control sounds not inducing RI in order to separate the neural signature of tinnitus suppression from tinnitus-unspecific neurophysiological effects.

## Acknowledgments

We are thankful to all participants for their time and patience and especially grateful to Susanne Staudinger for her help with the study management. Moreover, we thank Anita Hafner and Bernhard Unsin for their valuable support in data acquisition and study management.

## 8 Appendices

**Table S1:** Tinnitus loudness per condition. WN = white noise; IBP = individualized bandpass filtered white noise; IBS = individualized bandstop filtered white noise; M = mean; SD = standard deviation; Md = median; Min = minimum; Max = maximum; T0 = immediately after stimulation offset

|  | Total |  |  |  | T0 |  |  |  |
| --- | --- | --- | --- | --- | --- | --- | --- | --- |
|  | M ± SD | Md | Min | Max | M ± SD | Md | Min | Max |
| **Total sample** |  |  |  |  |  |  |  |  |
| WN | 88.29 ± 19.26 | 100.00 | 20.00 | 110.00 | 88.00 ± 21.60 | 90.00 | 20.00 | 110.00 |
| IBP | 87.24 ± 17.73 | 90.00 | 20.00 | 110.00 | 86.67 ± 21.95 | 90.00 | 20.00 | 110.00 |
| IBS | 92.60 ± 14.77 | 100.00 | 50.00 | 110.00 | 93.11 ± 16.35 | 100.00 | 50.00 | 110.00 |

**Table S2:**
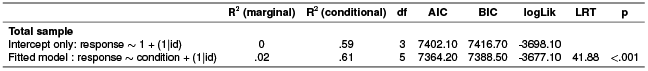
Model fitting - tinnitus loudness ratings. df = degrees of freedom; AIC = Akaike Information Criterion; BIC = Bayesian Information Criterion; logLik = log-likelihood; LRT = Likelihood Ratio Test

**Table S3:** Fixed effect testing - tinnitus loudness ratings. numDF = degrees of freedom numerator; denDF = degrees of freedom denominator

|  | numDF | denDF | F | p |
| --- | --- | --- | --- | --- |
| Condition | 2.00 | 900.00 | 21.43 | <.001 |

**Table S4:** Stimulus evaluation. WN = white noise; IBP = individualized bandpass filtered white noise; IBS = individualized bandstop filtered white noise; M = mean; SD = standard deviation; Md = median; Min = minimum; Max = maximum

|  | Valence |  |  |  | Arousal |  |  |  |
| --- | --- | --- | --- | --- | --- | --- | --- | --- |
|  | M ± SD | Md | Min | Max | M ± SD | Md | Min | Max |
| WN | .18 ± 1.83 | 0 | -4.00 | 4.00 | -.16 ± 1.68 | 0 | -4.00 | 3.00 |
| IBP | -.40 ± 2.18 | 0 | -4.00 | 4.00 | .51 ± 1.73 | 0 | -3.00 | 3.00 |
| IBS | 1.02 ± 2.09 | 2.00 | -4.00 | 4.00 | -.84 ± 1.82 | 0 | -4.00 | 3.00 |
| **Male** |  |  |  |  |  |  |  |  |
| WN | .03 ± 1.47 | 0 | -4.00 | 3.00 | .19 ± 1.25 | 0 | -3.00 | 3.00 |
| IBP | -.39 ± 2.26 | 0 | -4.00 | 4.00 | .68 ± 1.58 | 0 | -2.00 | 3.00 |
| IBS | .87 ± 1.98 | 1.00 | -4.00 | 4.00 | -.52 ± 1.69 | 0 | -4.00 | 3.00 |
| **Female** |  |  |  |  |  |  |  |  |
| WN | .50 ± 2.47 | 0 | -4.00 | 4.00 | .50 ± 2.47 | 0 | -4.00 | 4.00 |
| IBP | -.43 ± 2.06 | 0 | -4.00 | 3.00 | -.43 ± 2.06 | 0 | -4.00 | 3.00 |
| IBS | 1.36 ± 2.37 | 2.00 | -4.00 | 4.00 | -1.57 ± 1.95 | -2.00 | -4.00 | 2.00 |

**Table S5:**
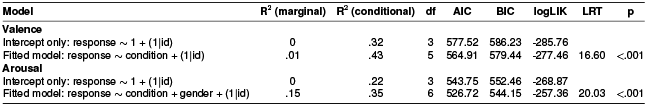
Model fitting - valence & arousal ratings. df = degrees of freedom; AIC = Akaike Information Criterion; BIC = Bayesian Information Criterion; logLik = log-likelihood; LRT = Likelihood Ratio Test

**Table S6:**
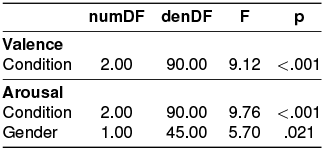
Fixed effect testing - valence & arousal ratings. numDF = degrees of freedom numerator; denDF = degrees of freedom denominator

**Table S7:** Post-hoc tukey contrasts for condition - stimulus evaluation. WN = white noise; IBP = individualized bandpass filtered white noise; IBS = individualized bandstop filtered white noise. Valence: Degrees of freedom = 92.00; standard error = .34; Arousal: Degrees of freedom = 92.00; standard error = .31; Gender: Degrees of freedom = 47.10; standard error = .39

| **Contrast** | **Valence** |  |  |  | **Arousal** |  |  |  |
| --- | --- | --- | --- | --- | --- | --- | --- | --- |
|  | **Estimate** | **t** | **p** | **d** | **Estimate** | **t** | **p** | **d** |
| WN - IBP | 0.58 | 1.71 | .209 | .288 | -.76 | -2.15 | .086 | .392 |
| **WN - IBS** | **-.84** | **-2.49** | **.038** | **.428** | 0.69 | 2.22 | .073 | .388 |
| **IBP - IBS** | **-1.42** | **-4.20** | **<.001** | **.665** | **1.36** | **4.37** | **<.001** | **.760** |
| **Male - Female** |  |  |  |  | **.90** | **2.33** | **.024** | **.728** |

**Table S8:**
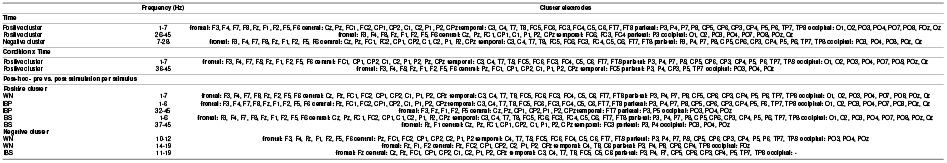
Cluster electrodes - total sample analysis. WN = white noise; IBP = individualized bandpass filtered white noise; IBS = individualized bandstop filtered white noise. Electrodes within frequency clusters grouped by brain areas.

**Table S9:**
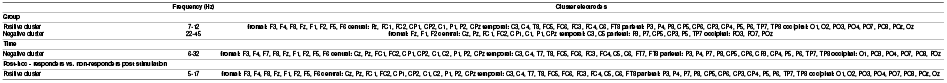
Cluster electrodes - non-responder vs. responder analysis. Electrodes within frequency clusters grouped by brain areas.

**Figure S1:**
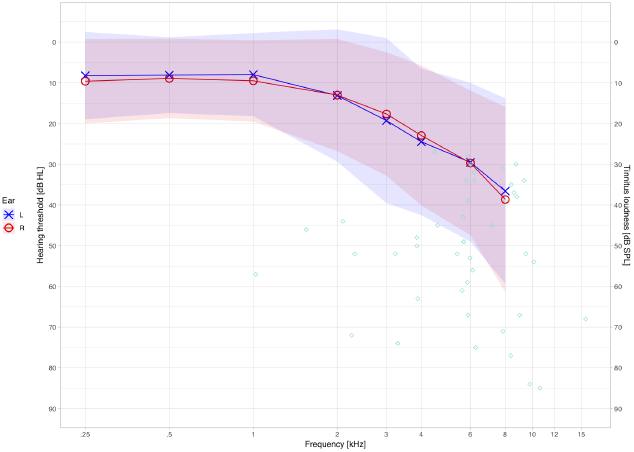
Audiometry and Tinnitometry. L = left; R = right; HL = hearing loss; SPL = sound pressure level. Results of audiometric assessment for both ears together with tinnitus frequency and loudness. The frequencies of hearing loss overlap with tinnitus frequencies.

**Figure S2:**
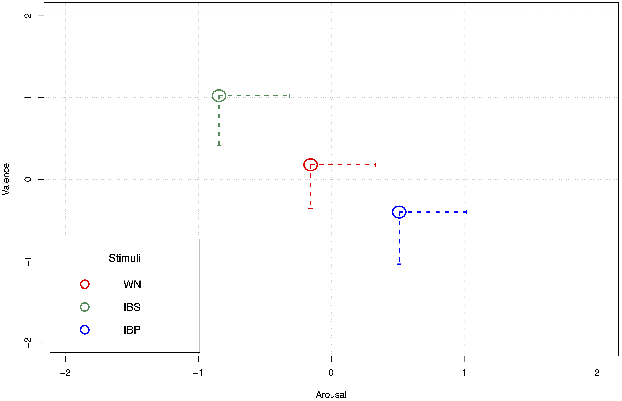
Valence and arousal evaluation per stimuli. WN = white noise; IBP = individualized bandpass filtered white noise; IBS = individualized bandstop filtered white noise. The value 0 indicates a neutral stimuli evaluation (cf. section acoustic stimulation). Highest tolerability was found for stimulus IBS as exemplified by high valence and low arousal ratings. While stimulus IBP resulted in the lowest tolerability evaluation. Parentheses display 95% confidence intervals for valence and arousal evaluation of the three stimuli.

